# Contrasting controls on tree methane emissions in upland and wetland forests

**DOI:** 10.64898/2026.01.29.702553

**Authors:** Jonathan Gewirtzman, Naomi Hegwood, Hannah Burrows, Maxwell Lutz, Grace Thompson, Bethany Duncan, Masako Yang, Sam Jurado, Robert Marra, Jaclyn Hatala Matthes

## Abstract

Trees can produce, consume, transport, and emit methane (CH₄), yet the environmental controls and mechanisms underlying these fluxes remain poorly understood. We combined 1,640 stem-chamber observations (2023–2025) with tower-based meteorology, soil moisture and temperature networks, water table monitoring, and non-destructive tomography to test how hydrology, energy balance, species identity, and internal wood condition regulate stem CH₄ flux. Wetland trees emitted approximately 40-fold more CH₄ than upland trees (1.96 vs. 0.05 nmol m⁻² s⁻¹). At the wetland, a three-way interaction between soil temperature, water table depth, and species explained 65% of flux variance, consistent with soil-derived CH₄ transport through stems. The wetland specialist *Nyssa sylvatica* emitted an order of magnitude more CH₄ than co-occurring generalists, likely reflecting flood-tolerance adaptations that enhance gas transport. In contrast, upland fluxes showed minimal environmental control (R² < 9%), with most variance occurring as unexplained temporal variation within individual trees—a pattern suggesting competing methanogenic and methanotrophic processes operating near equilibrium. Internal wood condition, assessed via acoustic and electrical resistance tomography, had opposite effects across sites: decay increased emissions in upland trees, likely by creating anaerobic microsites for in situ production, while decay decreased net emissions in wetland trees, likely by impairing transport of soil-derived CH₄ more than it enhanced in situ production. Together, these results indicate that the dominant controls on tree CH₄ flux differ fundamentally between wetland and upland forests, underscoring the need to represent hydrologic setting, species composition, and tree condition when scaling forest CH₄ contributions to regional budgets.

## Introduction

Forests can act as both CH₄ sources and sinks through complex biotic and abiotic pathways, but the magnitude and environmental controls on these fluxes remain poorly quantified (Covey and Megonigal 2019; Nazaries et al. 2013). Major discrepancies persist between top-down atmospheric models and bottom-up process-based inventories, with differences of nearly an order of magnitude (Saunois et al. 2020). Understanding CH₄ fluxes from underrepresented sources such as trees is essential for reducing these uncertainties and improving global CH₄ budgets.

Historically, research on forest CH₄ fluxes emphasized soil processes, where microbial methanogenesis in anoxic conditions produces CH₄ and methanotrophy in oxic conditions consumes it (Le Mer and Roger 2001; Bowden et al. 1998). However, trees are increasingly recognized as significant contributors, capable of emitting CH₄ through in situ production within tree tissues, transport from soil via the vascular system, and diffusion through tree surfaces (Pangala et al. 2017; Yip et al. 2019). Wetland ecosystems are generally net methane producers (Delwiche et al. 2021), and trees in these systems transport and emit soil-derived CH₄ directly to the atmosphere, bypassing oxidation in the soil and contributing substantially to ecosystem budgets (Barba et al. 2021; Jeffrey et al. 2023). Upland ecosystems are generally net methanotrophic (Le Mer and Roger 2001; Conrad 2009) but the extent to which tree fluxes offset ecosystem uptake remains highly uncertain (Pitz and Megonigal 2017; Gauci et al. 2024).

The microbiome of tree wood may play a central role in stem CH₄ dynamics (Baldrian 2017; Mishra et al. 2020; Uroz et al. 2016; Arnold et al. 2025). Heartwood and wetwood—waterlogged, often degraded internal wood—harbor methanogenic archaeal communities adapted to low-oxygen conditions within living trees (Schink et al. 1981; Zeikus and Ward 1974; Gewirtzman et al. 2025; Yip et al. 2019; Putkinen et al. 2021; Feng et al. 2022; Harada et al. 2024). Internal decay may therefore create anaerobic microsites favorable for in situ CH₄ production, while also potentially altering gas transport pathways. Meanwhile, both upland and wetland trees may exhibit methanotrophic activity at some woody surfaces (Gauci et al. 2024).

Tree-mediated CH₄ fluxes also vary with environmental conditions and species-specific traits including wood density and secondary metabolites (Moisan et al. 2024). Fluxes vary substantially among species and along with tree diameter, temperature, and soil moisture, with emissions often exhibiting threshold responses to volumetric water content (Epron et al. 2022; Hettwer et al. 2025; Barba et al. 2024).

This study focuses on tree-mediated CH₄ fluxes at Harvard Forest, a long-term ecological research site in north-central Massachusetts that includes a ∼14,000-year-old peatland and adjacent upland forests. We measured two generalist species, red maple (*Acer rubrum*) and eastern hemlock (*Tsuga canadensis*), and one specialist species at each site: black gum (*Nyssa sylvatica*) in the wetland and red oak (*Quercus rubra*) in the upland, repeatedly across seasons for two years.

We address three questions. First, how do stem CH₄ fluxes vary seasonally and across a wetland–upland gradient, and to what extent are fluxes regulated by meteorological and hydrologic drivers? Second, how do species modulate the sensitivity of stem CH₄ fluxes to hydrological and energy drivers, and do individual trees exhibit consistent flux behavior through time? Third, does internal wood condition, including decay and wetwood, modify tree CH₄ emissions? By integrating repeated stem-chamber measurements with tower-based meteorological data and hydrologic observations, we quantify how vegetation mediates methane exchange under variable climate conditions.

## Methods

### Study site

This study was conducted at Harvard Forest in north-central Massachusetts, USA (42.5°N, 72.2°W, elevation 220–410 m). The regional climate is cool and moist with precipitation distributed evenly throughout the year. Based on 1991–2020 meteorological data, mean annual temperature was 8.2°C and mean annual precipitation was 1175 mm (Boose and VanScoy 2025a). The study years 2023 and 2024 were slightly warmer and wetter than average, with mean annual temperatures of 9.2°C and 9.1°C and total precipitation of 1482 mm and 1384 mm, respectively (Boose and VanScoy 2025a).

Harvard Forest lies within the transition hardwood-white pine-hemlock region of the northeastern United States. Current stands are 75–125 years old, reflecting agricultural abandonment, extensive logging of white pine (Pinus strobus), and damage from the 1938 hurricane (Barker-Plotkin et al. 2015a). Dominant species include red oak, red maple, white pine, and eastern hemlock (Jenkins et al. 2008). Metamorphic bedrock underlies acidic, stony glacial tills interspersed with glacial outwash deposits in wetland basins (Anderson et al. 2003; Barker-Plotkin et al. 2015b). Upland soils are moderately to well-drained; basin and valley soils are poorly to excessively drained (Anderson et al. 2003; Jenkins and Motzkin 2023).

Measurements were conducted at two sites within the Prospect Hill tract representing contrasting hydrological regimes. The upland site was located within the footprint of the Environmental Measurements Site (EMS) eddy covariance tower, where 40 inventory plots (10 m radius) extend up to 500 m from the tower (J. Matthes et al. 2025); nine plots were used for tree selection. The wetland site was Black Gum Swamp (BGS), a groundwater-fed peat basin (∼200 × 600 m) approximately 1 km from EMS (Anderson et al. 2003). BGS is characterized by an abrupt transition from surrounding dry oak forest to lowland peat, with microtopography creating drier hummocks and wetter hollows.

### Tree selection and sampling design

We selected 60 trees for stem CH₄ flux measurements, 30 per site. The design included two habitat generalists present at both sites—eastern hemlock (*T. canadensis*) and red maple (*A. rubrum*)—and one specialist at each site: black gum (*N. sylvatica*) at the wetland and red oak (*Q. rubra*) at the upland. Ten individuals of each species were selected at each site where that species occurred. Trees were selected to represent typical sizes rather than external indicators of decay. Using diameter at breast height (DBH) measurements from the Harvard Forest ForestGEO plot, we defined the second and third quartiles of the DBH distribution for each species and selected trees within this range.

### Stem flux measurements

#### Collar installation

Permanent stem flux collars were installed on each tree in May 2023, positioned 1 m above the tree base. Collars were constructed from circular polyvinyl chloride (PVC) pipe (inner diameter 10.16 cm, depth 3.8–5.1 cm). Two collar types were used depending on stem curvature: curved collars (interior volume 457.39 cm³) with inner surfaces shaped to conform to the stem, and non-curved collars (544.39 cm³) for flatter stems. Collars were attached using non-VOC silicone sealant and held in place with elastic cord while curing for at least 24 hours.

#### Gas measurements

Measurements were made in situ using a portable Ultra-Portable Greenhouse Gas Analyzer (UGGA; Los Gatos Research, Mountain View, CA, USA), which uses off-axis integrated cavity output spectroscopy (OA-ICOS) to measure CH₄ and CO₂ at 1 Hz. During measurements, a cap lined with closed-cell foam was placed over the collar to create a closed chamber, with the foam gasket compressing against the collar rim. Gas circulated continuously through the analyzer and returned to the closed system. Both CO₂ and CH₄ were recorded for a minimum of 3 minutes per measurement.

CO₂ concentrations were monitored in real time to verify seal integrity. Prior to each measurement, collars were leak-tested by breathing around the collar-cap junction while the analyzer was attached; leaks produced a rapid CO₂ spike. When leaks were detected, additional sealant was applied and the collar was allowed to cure before re-measurement.

Measurements extended from June 2023 through October 2025. Frequency was adjusted seasonally: approximately weekly during summer (June–August), biweekly during spring (March–May) and fall (September–November), and monthly during winter (December–February).

### Flux calculations

Flux rates were calculated from the linear rate of change in gas concentration over time:

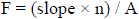

where F is flux (nmol m⁻² s⁻¹ for CH₄; µmol m⁻² s⁻¹ for CO₂), slope is from linear regression of concentration versus elapsed time, n is total moles of gas in the closed system, and A is bark surface area enclosed by the collar (assuming a cylindrical section with 10.16 cm diameter). Moles were calculated using the ideal gas law with atmospheric pressure and air temperature from the Fisher Meteorological Station (Boose and VanScoy 2025a). Collar depth was measured at four interior locations and averaged to calculate mean collar volume for each tree.

### Tomographic assessment of internal decay

We used acoustic and electrical resistance tomography to non-destructively assess internal wood condition in each tree. All measurements were conducted at breast height (1.37 m) using the PiCUS system (Argus Electronic GmbH, Rostock, Germany).

#### Sonic tomography

Sonic tomography (SoT) detects internal decay by measuring acoustic wave velocity through wood (Brazee et al. 2011). A minimum of 8 sensors were placed around the trunk circumference following manufacturer-recommended spacing. Sterilized steel nails were inserted through bark to contact wood, and sensor positions were recorded using the PiCUS Caliper 3, which determines cross-sectional geometry through triangulation. Each point was struck 3–5 times with an electronic hammer, and travel times were recorded between all sensor pairs.

The PiCUS Q74 software generates cross-sectional images of apparent sonic velocity via tomographic reconstruction. Decayed wood transmits sound more slowly due to reduced modulus of elasticity and density; the software compares measured travel times to expected times through solid wood within the same trunk, facilitating cross-species comparisons (Gilbert et al. 2016). Tomograms display brown (high velocity, sound wood), green (intermediate velocity, reduced density), and blue/violet (low velocity, advanced decay or cavity). We exported percent sound wood (brown pixels) and percent decay (non-brown pixels).

#### Electrical resistance tomography

Electrical resistance tomography (ERT) measures the spatial distribution of electrical resistance across the cross-section (Brazee et al. 2011). Resistance is influenced by moisture content, ion concentration, and cell structure: low resistance indicates elevated moisture or ions often associated with active fungal decay or bacterial wetwood; high resistance indicates dry wood or cavities. We used the TreeTronic 3 Tomograph, attaching electrode clamps to the same nails used for SoT. The system sequentially injects current between electrode pairs and measures voltage distribution, reconstructing spatial resistivity patterns (Ω·m).

The PiCUS software does not provide quantitative ERT summary statistics. We developed a custom image analysis application in R using the Shiny framework (Chang et al. 2026) to extract numeric metrics from exported images. The application extracts the colorbar, constructs a log-transformed calibration function relating pixel colors to resistivity, and applies this calibration to pixels within a user-delineated cross-sectional boundary. For each tree we calculated mean and median resistivity, standard deviation, and coefficient of variation (CV)..

#### Decay classification

Trees were classified into four decay categories based on joint interpretation of SoT and ERT results, following (Brazee et al. 2011) and manufacturer guidelines (Table 1). Thresholds were >1% damage from SoT (percent non-brown pixels) and ERT CV > 0.5.

**Table 1.**
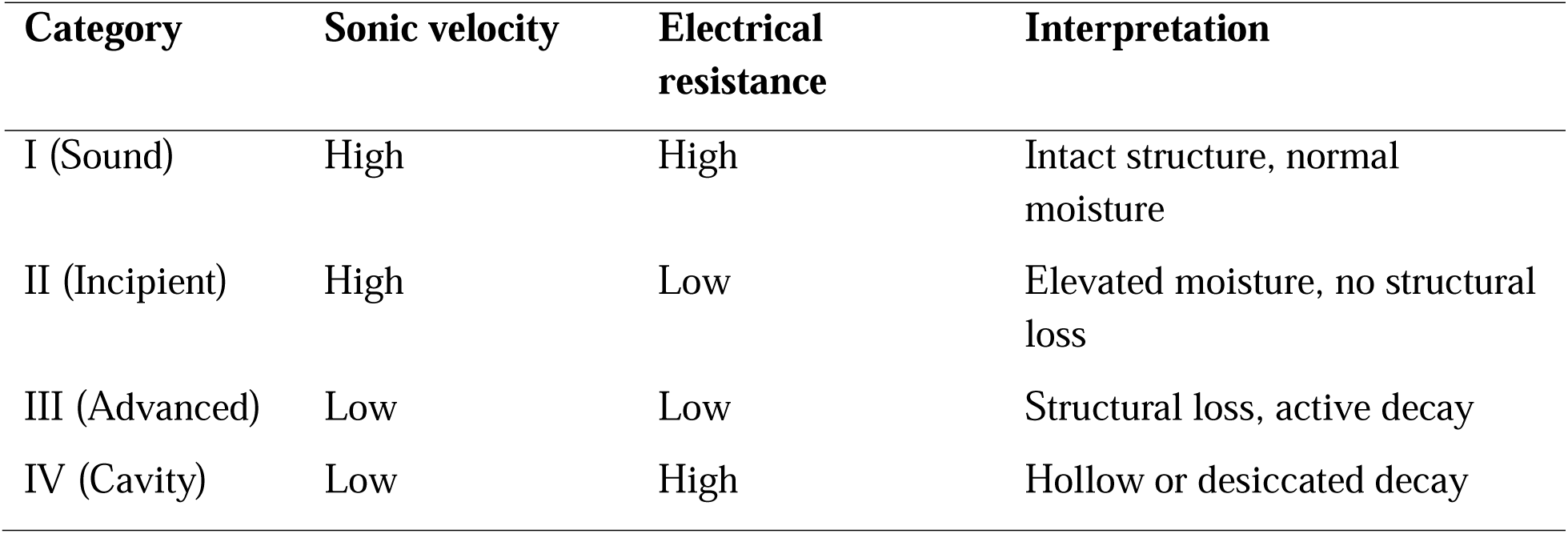
Decay classification system based on combined sonic tomography (SoT) and electrical resistance tomography (ERT) results.

### Environmental data

Environmental data were compiled from multiple monitoring networks at Harvard Forest. From the Fisher Meteorological Station (Boose and VanScoy 2025a), we obtained 15-minute measurements of air temperature, relative humidity, atmospheric pressure, precipitation, photosynthetically active radiation (PAR), net radiation, global solar radiation, and soil temperature at 10 cm depth. Vapor pressure deficit (VPD) was calculated from air temperature and relative humidity.

Ecosystem-scale fluxes were obtained from the EMS eddy covariance tower (Munger and Wofsy 2024): hourly net ecosystem exchange (NEE) of CO₂, latent heat flux (LE), sensible heat flux (H), and friction velocity (u*). Understory microclimate data came from the EMS understory network (Munger and Hadley 2025): soil temperature at 10 cm and volumetric water content in the upper 15 cm. Water table depth was measured at 15-minute intervals at two wetland stations (Boose and VanScoy 2025b): Black Gum Swamp (BGS) and Beaver Swamp (BVS).

Local soil volumetric water content and temperature were measured at three locations around each study tree using ProCheck handheld readers with GS3 sensors (METER Group, Pullman, WA, USA). Soil water content at multiple depths was obtained from the NEON Terrestrial Instrument System (National Ecological Observatory Network (NEON) 2026) and summarized to 30-minute averages from shallow (∼6 cm), mid (∼16–26 cm), and deep (∼50+ cm) sensors. Canopy phenology metrics (green chromatic coordinate and NDVI) were derived from the PhenoCam network camera at EMS (Seyednasrollah et al. 2019; Richardson et al. 2018). All time series were aligned to hourly resolution.

### Statistical analysis

#### Rolling window correlation analysis

To identify temporal scales at which environmental variables most strongly correlate with stem CH₄ flux, we calculated right-aligned rolling means for each predictor across window sizes in 3 hour intervals to 14 days. For each window size, we calculated Pearson correlations between the rolling mean and instantaneous CH₄ flux, separately by site. For each predictor, we identified the window yielding the strongest correlation (by absolute value) and applied Benjamini-Hochberg false discovery rate correction (FDR < 0.05) across predictors within each site.

#### Model development

We used hierarchical clustering of the correlation matrix among significant predictors to identify redundant variables (|r| ≥ 0.7), retaining the predictor with the strongest CH₄ correlation and lowest missing data from each cluster. Based on prior ecological knowledge, we designated soil temperature and a hydrological variable (water table depth for wetland, soil water content for upland) as core predictors for all models. Additional predictors were selected via randomized forward selection: in each of 100 iterations, candidates were added in random order, retaining those that improved AIC until no further improvement or a maximum of 6 predictors. Predictors appearing in >50% of iterations were included in final models.

We fit linear mixed-effects models using lme4 in R (Bates et al. 2015). To accommodate right-skewed distributions and negative fluxes, we applied an inverse hyperbolic sine (asinh) transformation after scaling by 1000. The asinh transformation approximates natural log for large values while remaining defined for zero and negative values. All continuous predictors were standardized prior to fitting.

We tested all two-way and three-way interactions among core predictors and species. Final models included a three-way interaction between soil temperature, hydrological status, and species, with individual tree as a random intercept. Separate models were fit for each site. Model fit was assessed using R², and variance inflation factors (VIF) were calculated. Predictions were back-transformed using sinh for visualization.

#### Tree-level repeatability

To assess whether individual trees exhibited consistent CH₄ flux behavior over time, we calculated the intraclass correlation coefficient (ICC) for each species × site combination from an intercept-only mixed model with tree as random effect. We divided the study at June 1, 2024 into early and late periods, calculated each tree’s mean flux in each period, and computed Spearman rank correlations between periods within each group. Likelihood ratio tests compared models with and without tree random effects.

### Software and data availability

Analyses were conducted in R version 4.3.x (R Core Team 2023). Key packages included lme4 (Bates et al. 2015), emmeans, RcppRoll, shiny (Chang et al. 2025), and tidyverse (Wickham et al. 2019). Environmental data are available through the Harvard Forest Data Archive (https://harvardforest.fas.harvard.edu/data), NEON (https://data.neonscience.org), and PhenoCam (https://phenocam.nau.edu). Code will be available on Github, and flux data on the Harvard Forest Data Archive, upon publication.

## Results

### Temporal and spatial patterns in stem CH₄ flux

We measured stem CH₄ flux from 60 trees (30 per site, 10 per species) across 29 sampling rounds from June 2023 to October 2025, yielding 1,640 observations after quality control (Figure 1). Stem fluxes were substantially higher at the wetland site (mean 1.96 ± 0.27 nmol m⁻² s⁻¹, median 0.15, IQR 0.04–0.59) than the upland site (mean 0.05 ± 0.01 nmol m⁻² s⁻¹, median 0.02, IQR −0.003–0.08), representing an approximately 40-fold difference in mean emission rates. Positive CH₄ emissions were observed in 73.5% of upland measurements and 90% of wetland measurements.

**Figure 1.**
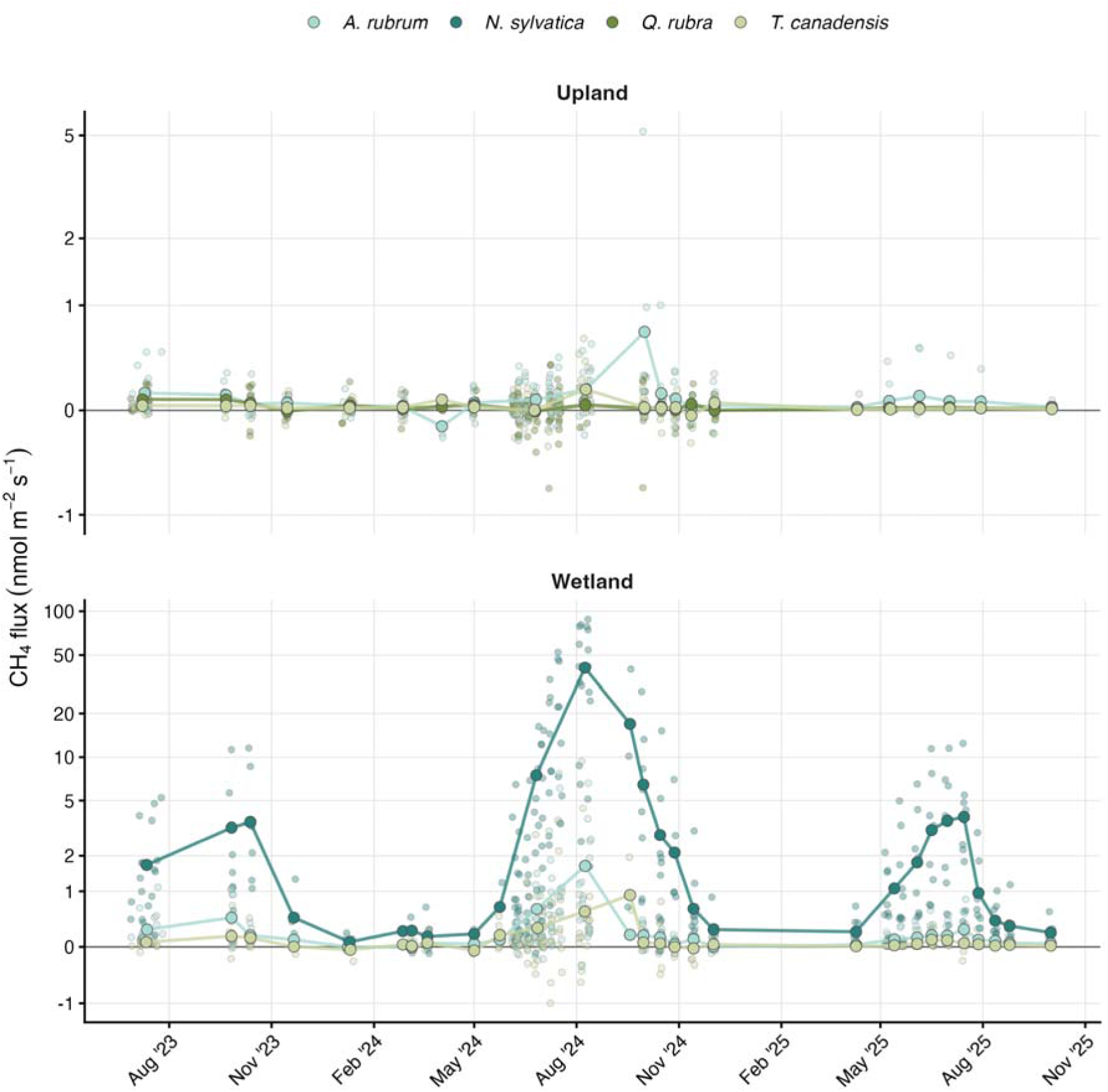
Stem methane flux over time at wetland (top) and upland (bottom) sites. Individual observations (transparent points) and sampling round means (connected points) are shown for each species. Sampling rounds were defined as measurement campaigns separated by >7 days. Y-axis is asinh-transformed to display both positive emissions and negative fluxes (uptake).

Species identity strongly influenced flux magnitude, particularly at the wetland site (Figure 2). Nyssa sylvatica exhibited the highest emissions (tree-level mean 5.67 ± 1.61 nmol m⁻² s⁻¹, median 4.17), exceeding the next highest species by more than an order of magnitude. At the wetland site, N. sylvatica emitted significantly more CH₄ than both Acer rubrum (difference = 5.29 nmol m⁻² s⁻¹, p = 0.001) and Tsuga canadensis (difference = 5.48 nmol m⁻² s⁻¹, p < 0.001), which did not differ from each other (p = 0.99). At the upland site, A. rubrum showed higher emissions than both Quercus rubra (p = 0.027) and T. canadensis (p = 0.036), while these latter two species did not differ (p = 0.99). The habitat generalist A. rubrum showed significantly higher emissions at the wetland than upland site (3.4-fold, p = 0.009). T. canadensis showed a similar but non-significant pattern (6.3-fold, p = 0.16).

**Figure 2.**
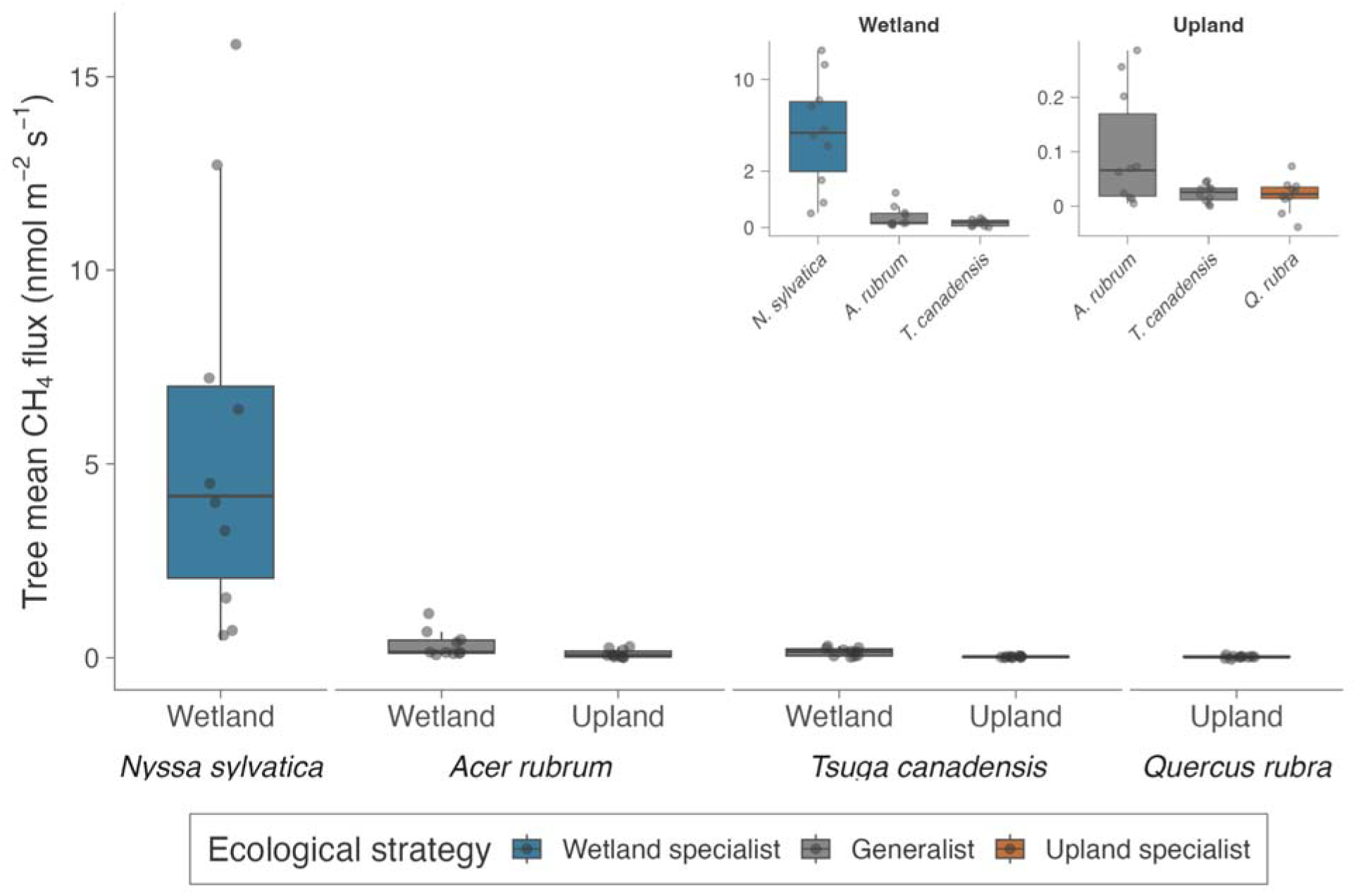
Tree mean CH₄ flux by species and site. Main panel shows boxplots of tree-level mean fluxes (n = 10 trees per species × site combination) with individual trees as points. Species are grouped by ecological strategy: wetland specialist (*N. sylvatica*), upland specialist (*Q. rubra*), and habitat generalists (*A. rubrum*, *T. canadensis*). Inset panels show the same data on pseudo-log scales to visualize differences among lower-emitting groups.

Stem CH₄ fluxes exhibited strong seasonality at both sites, with peak emissions in August (wetland: 7.04 ± 1.80 nmol m⁻² s⁻¹; upland: 0.14 ± 0.03 nmol m⁻² s⁻¹) and minimum emissions outside of the growing season (wetland January: 0.01; upland April: 0.003 nmol m⁻² s⁻¹). Summer 2024 showed notably elevated emissions at the wetland site compared to other years (mean 5.30 ± 0.90 vs. 0.78 ± 0.18 in 2023 and 0.75 ± 0.13 nmol m⁻² s⁻¹ in 2025), driven almost entirely by N. sylvatica (summer mean 14.8 in 2024 vs. 1.7 in 2023 and 2.08 nmol m⁻² s⁻¹ in 2025). Upland fluxes showed minimal interannual variation.

### Tree-level repeatability

Intraclass correlation coefficients (ICC) were low across all species × site combinations (range 0–0.11, mean 0.05), indicating that between-tree variance accounted for only a small fraction of total variance in CH₄ flux (Figure 3). N. sylvatica at the wetland site showed the highest ICC (0.108, LRT p < 0.001), followed by A. rubrum at the wetland (0.104, p < 0.001) and upland (0.052, p = 0.056) sites. T. canadensis at the upland site showed essentially zero repeatability (ICC = 0, LRT p = 1.0).

**Figure 3.**
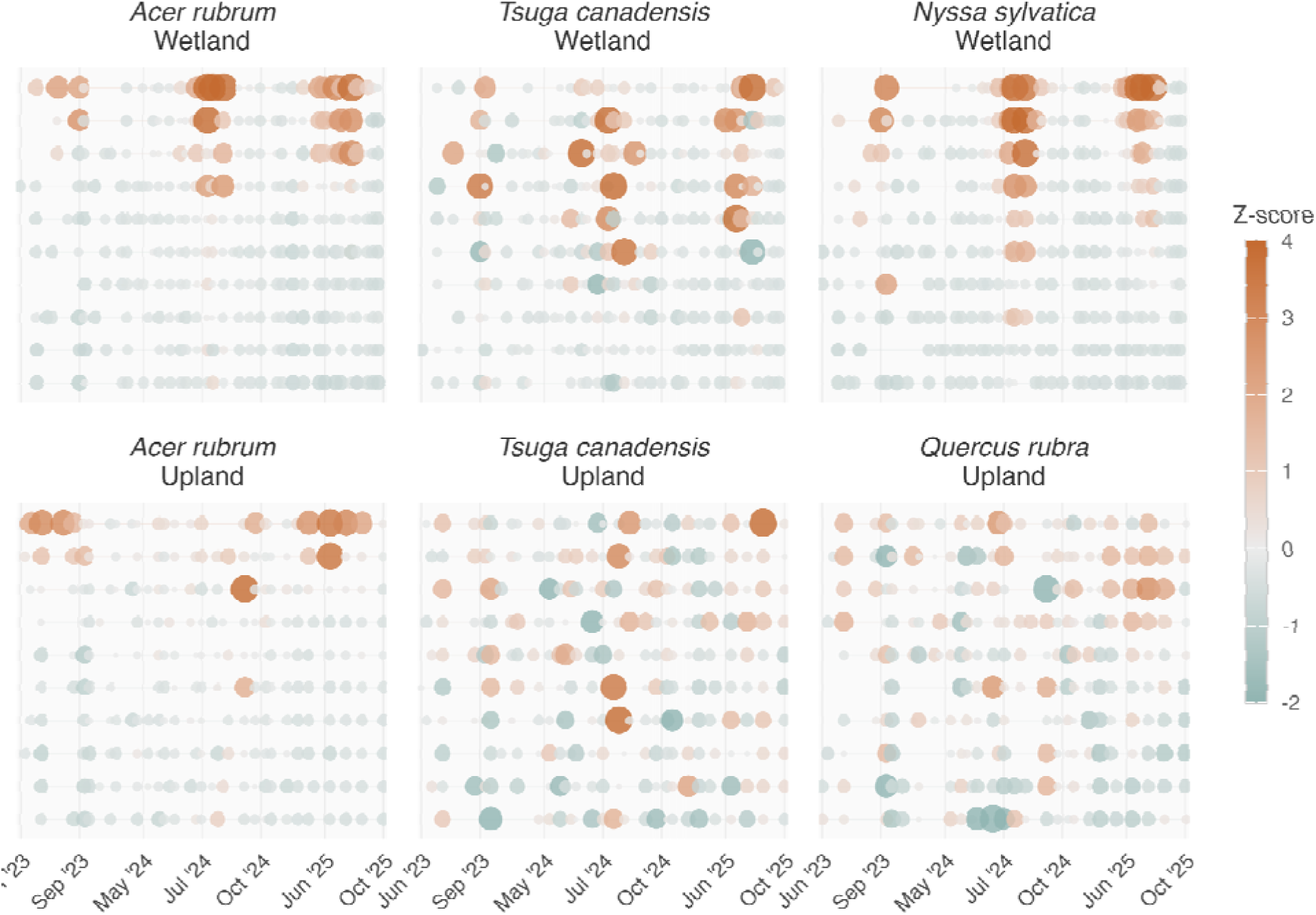
(z-score tracks). Individual tree flux trajectories shown as z-scores (standardized within year × site × species). Each row represents one tree, ordered from highest (top) to lowest (bottom) mean z-score. Point size and color indicate deviation from group mean. Trees maintain relatively consistent ranks over time despite substantial temporal variation in absolute flux.

Despite low ICCs, Spearman rank correlations between early-period (before June 2024) and late-period (after June 2024) tree means were consistently positive and moderate to strong (ρ = 0.56–0.87). N. sylvatica showed the strongest temporal consistency (ρ = 0.87, p = 0.003), followed by A. rubrum at the upland site (ρ = 0.75, p = 0.018). Trees maintained their relative rankings over time despite substantial variation in absolute flux.

### Environmental drivers

Rolling-window correlation analysis identified distinct environmental drivers at the two sites, with the wetland showing far more environmental sensitivity than the upland (26 vs. 5 significant predictors after FDR correction; Figure 4). At the wetland site, the strongest correlations were with atmospheric CO₂ concentration (r = −0.26 at 11.25d window), sensible heat flux (r = −0.23 at 5.9d), ecosystem CO₂ flux (r = −0.21 at 9h), precipitation (r = +0.21 at 8.25d), and shallow soil water content (r = +0.21 at 2.6d). Water table depth showed strongest correlations at short timescales (r = +0.18–0.20 at 18-h windows for both wetland monitoring wells).

**Figure 4.**
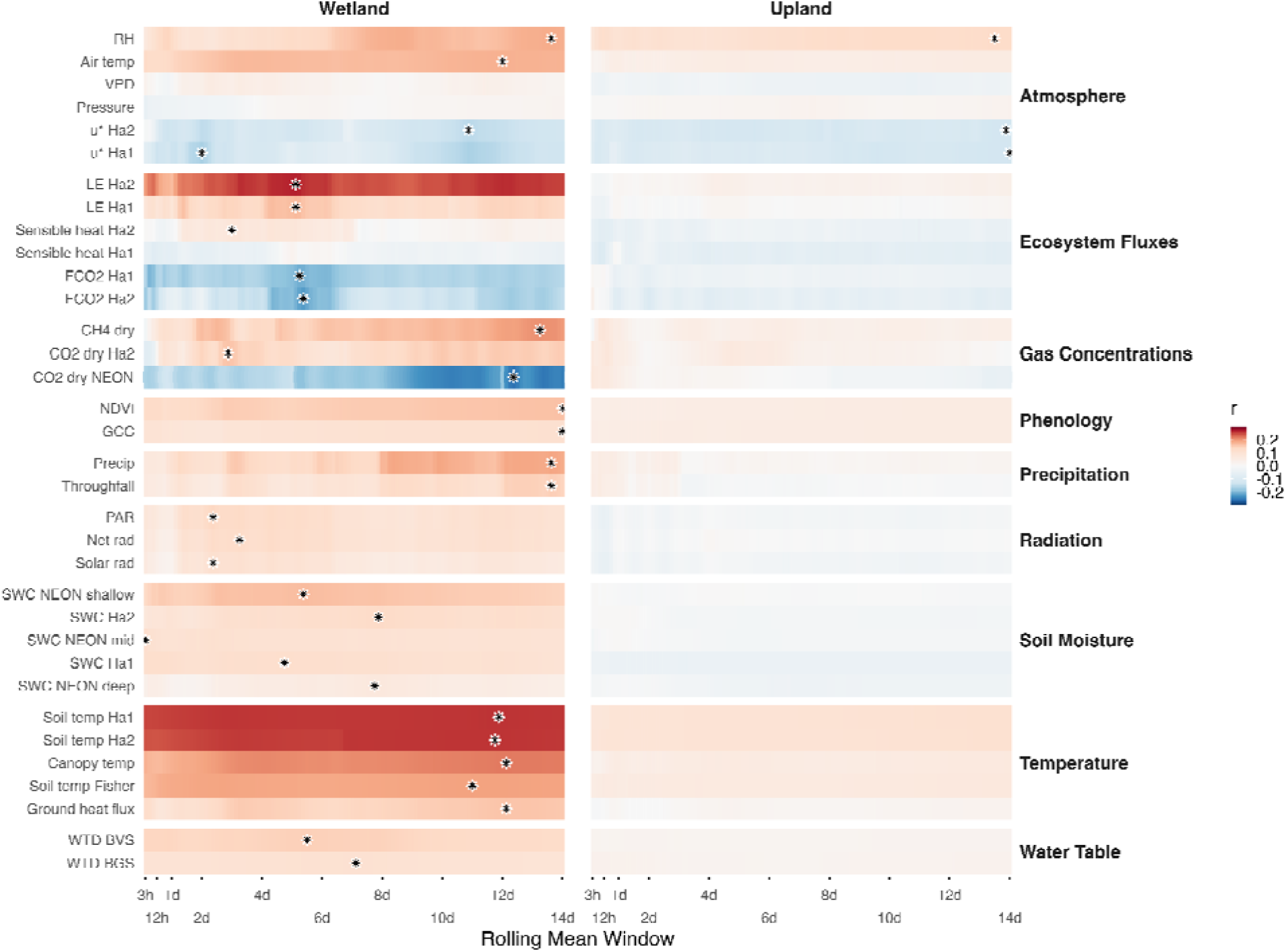
(rolling correlations). Pearson correlations between stem CH₄ flux and environmental variables across rolling mean windows from 3 hours to 14 days. Each row shows one predictor variable, grouped by category. Color indicates correlation strength and direction (blue = negative, red = positive) as a function of averaging timescale (x-axis). Asterisks indicate significant correlations after Benjamini-Hochberg FDR correction (α = 0.05) at the optimal window. Variables are ordered within each group by strongest absolute correlation.

Positive correlations with moisture-related variables (soil water content, water table, precipitation, relative humidity, latent heat flux) and negative correlations with variables associated with dry conditions (VPD, sensible heat flux, CO₂ concentration) indicate that wetter conditions enhance stem CH₄ emissions. Soil temperature was positively correlated with emissions at the wetland site (r = 0.07–0.14 at 1–3 d windows). Optimal averaging windows differed between sites: upland correlations were strongest at longer timescales (median 264 h, range 261–336 h), while wetland correlations spanned a broader range with a shorter median (57 h, range 3–291 h).

Five environmental variables were significantly correlated with CH₄ flux at both sites: water table depth (BGS well), VPD, solar radiation, relative humidity, and net radiation. All showed consistent directions of effect across sites. Twenty-one additional variables were significant only at the wetland site, including all soil moisture measurements, precipitation, throughfall, ecosystem fluxes (LE, FCO₂, sensible heat), atmospheric CO₂ concentrations, and soil temperature. No variables were significant only at the upland site.

### Wetland environmental model

A core model including soil temperature (282-h window), water table depth (132-h window), species, and all two-way and three-way interactions explained 65.4% of variance. All main effects and interactions were significant (p < 0.01). Species differed substantially in their temperature responses: N. sylvatica showed the strongest effect (standardized effect = 0.552), while T. canadensis (0.078) and A. rubrum (0.111) showed weaker responses. Water table effects followed a similar pattern (N. sylvatica: 0.534; T. canadensis: 0.074; A. rubrum: 0.190). The temperature × water table interaction was strongest for N. sylvatica (0.415) and weakest for T. canadensis (0.038), with A. rubrum intermediate (0.165). The intraclass correlation coefficient was 0.47, indicating substantial tree-to-tree variation after accounting for environmental predictors.

Alternative models substituting soil water content or latent heat flux for temperature or water table performed somewhat worse (Table SX), confirming temperature and water table as the primary drivers. A full model adding latent heat flux and shallow soil water content to the core model improved fit modestly compared to the base core model (R² = 0.676, AIC = 784.8), but the effects of these additional predictors were not robust: both reversed sign when controlling for temperature and water table, indicating they captured residual covariance rather than independent mechanisms. The core temperature × water table × species interactions remained similar between models.

### Upland environmental model

We developed analogous models for the upland site using 570 observations from 30 trees from June 2023 to April 2025. We tested two formulations: Model A used instantaneous (1-h window) soil temperature and soil water content; Model B used the same predictors and windows as the wetland model to enable direct comparison.

Both models explained minimal variance (Model A: R² = 8.8%; Model B: R² = 8.2%), and the models were statistically equivalent (ΔAIC = 57). Intraclass correlations were low (0.067), indicating most residual variance was temporal rather than between-tree. Species-specific responses differed markedly from the wetland: only A. rubrum showed significant environmental effects (positive temperature response, 0.056, p = 0.005), while T. canadensis and Q. rubra showed no significant responses to any predictor.

### Internal wood condition and stem CH₄ flux

We characterized internal wood condition in all 60 trees using electrical resistivity tomography (ERT) and sonic tomography. Trees were classified into four decay categories: Category I (intact; 50% of trees), Category II (low resistance only; 26.7%), Category III (both low resistance and structural damage; 15%), and Category IV (structural damage only; 8.3%). Mean ERT-measured moisture CV was similar between sites (wetland: 0.522 ± 0.192; upland: 0.526 ± 0.249), but upland trees showed more extensive SoT-measured structural damage (6.6 ± 10.6% damaged area) compared to wetland trees (1.3 ± 4.7%), with 36.7% of upland trees vs. 10% of wetland trees exceeding the 1% damage threshold.

The relationship between internal wood condition and CH₄ flux (Figure 5) differed between sites and was opposite in direction. At the upland site, ERT CV was positively correlated with flux (r = 0.597, p < 0.001), driven by significant positive correlations in Q. rubra (r = 0.802, p = 0.005) and A. rubrum (r = 0.665, p = 0.036). At the wetland site, ERT CV was negatively correlated with flux (r = −0.404, p = 0.027), driven by N. sylvatica (r = −0.68, p = 0.031). T. canadensis showed no significant relationship at either site.

**Figure 5.**
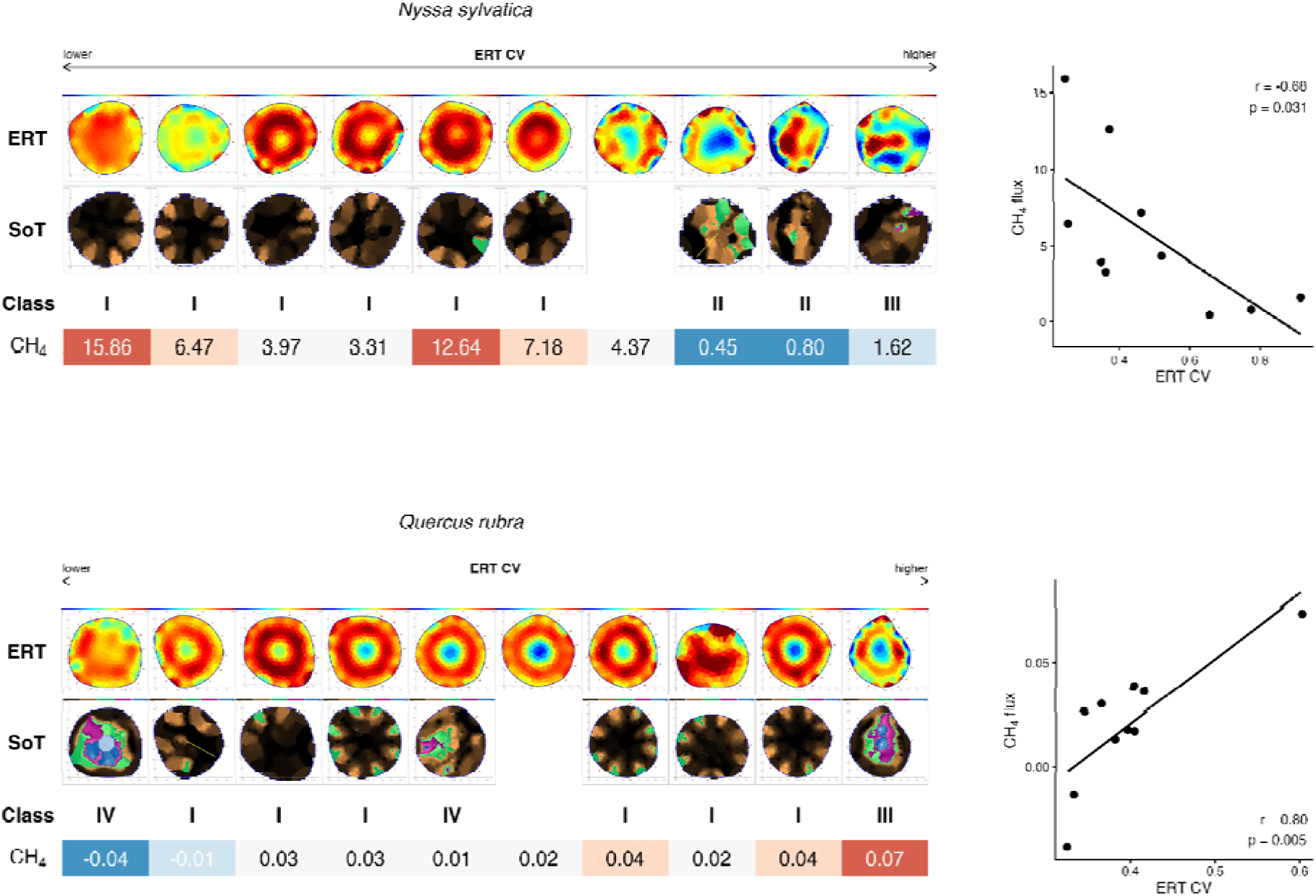
Internal wood condition and stem CH₄ flux across species and sites. For each species/location group, trees are arranged by increasing ERT coefficient of variation (CV; arrow at top). ERT: electrical resistivity tomography cross-sections showing internal electrical resistance patterns (blue = low resistance = wet, red = high resistance = dry). SoT: sonic tomography cross-sections showing wood density (blue/purple = low density/damaged, green = high density/sound). Class: decay category (I = intact; II = low resistance only; III = low resistance + high damage; IV = high damage only). CH₄: mean stem flux (nmol m⁻² s⁻¹), color-scaled within species (blue = low, red = high). Scatterplots show ERT CV vs CH₄ flux with correlation statistics. Upland species including *Q. rubra* show positive correlations between decay and flux, while wetland *N. sylvatica* shows negative correlation. Two SoT scans were incomplete and are not displayed.

### Variance partitioning

To quantify sources of variability, we partitioned variance using nested random effects models. When examining individual observations, site explained 21.4% of total variance, with the remaining variance reflecting within-site heterogeneity—particularly at the wetland where variance was 1,200-fold higher than upland. Within-site variance structure differed between locations. At the wetland, species identity explained 44.5% of variance, individual tree effects explained 18.5%, and residual variation explained 54.8% (ICC = 0.535). At the upland, residual variation dominated (86.3%), with minimal contributions from species identity (4.9%) or individual tree effects (10.2%; ICC = 0.149).

## Discussion

The drivers and mechanisms of stem CH₄ flux differed fundamentally between wetland and upland forests. At the wetland, temperature and water table interactions explained most variance, with species identity and individual tree effects accounting for substantial additional variation. At the upland, environmental predictors explained little variance, with most variation remaining unexplained and likely reflecting competing methanogenesis and methanotrophy operating near equilibrium. Internal wood condition showed opposite relationships with flux across sites: decay increased emissions at the upland but decreased emissions at the wetland, reflecting fundamentally different CH₄ sources and transport pathways.

### Wetlands: soil-derived CH₄ transport dominates

The strong temperature × water table interaction at the wetland site reflects synergistic effects of substrate availability, microbial activity, and transport physics on stem CH₄ emissions (Terazawa et al.; Christensen et al. 2003, Segers et al. 1998, Dunfield 1993). Figure 6A illustrates this synergy: under dry conditions (10th percentile water table), predicted flux remained near zero (0.14–0.43 nmol m⁻² s⁻¹) across the full temperature range, while under wet conditions (90th percentile), flux increased from 0.02 to 5.69 nmol m⁻² s⁻¹ across the same gradient. This pattern is consistent with a mechanistic framework where elevated water tables create anoxic conditions necessary for methanogenesis, while temperature controls methanogen growth rate, reaction kinetics, and potentially substrate supply through enhanced root exudation or organic matter decomposition. The temporal pattern of observed emissions supports this interpretation: periods when both soil temperature and water table were elevated (shaded regions in Figure 6B) corresponded with or preceded major flux peaks, including the pronounced summer 2024 emissions (peak 13.9 nmol m⁻² s⁻¹). In contrast, the upland site showed minimal environmental control (maximum predicted flux 0.079 nmol m⁻² s⁻¹), suggesting that factors not captured by tower-based or well-based measurements, such as tree-specific hydraulic properties, internal substrate dynamics, or localized microsite conditions, dominated upland flux variability.

**Figure 6.**
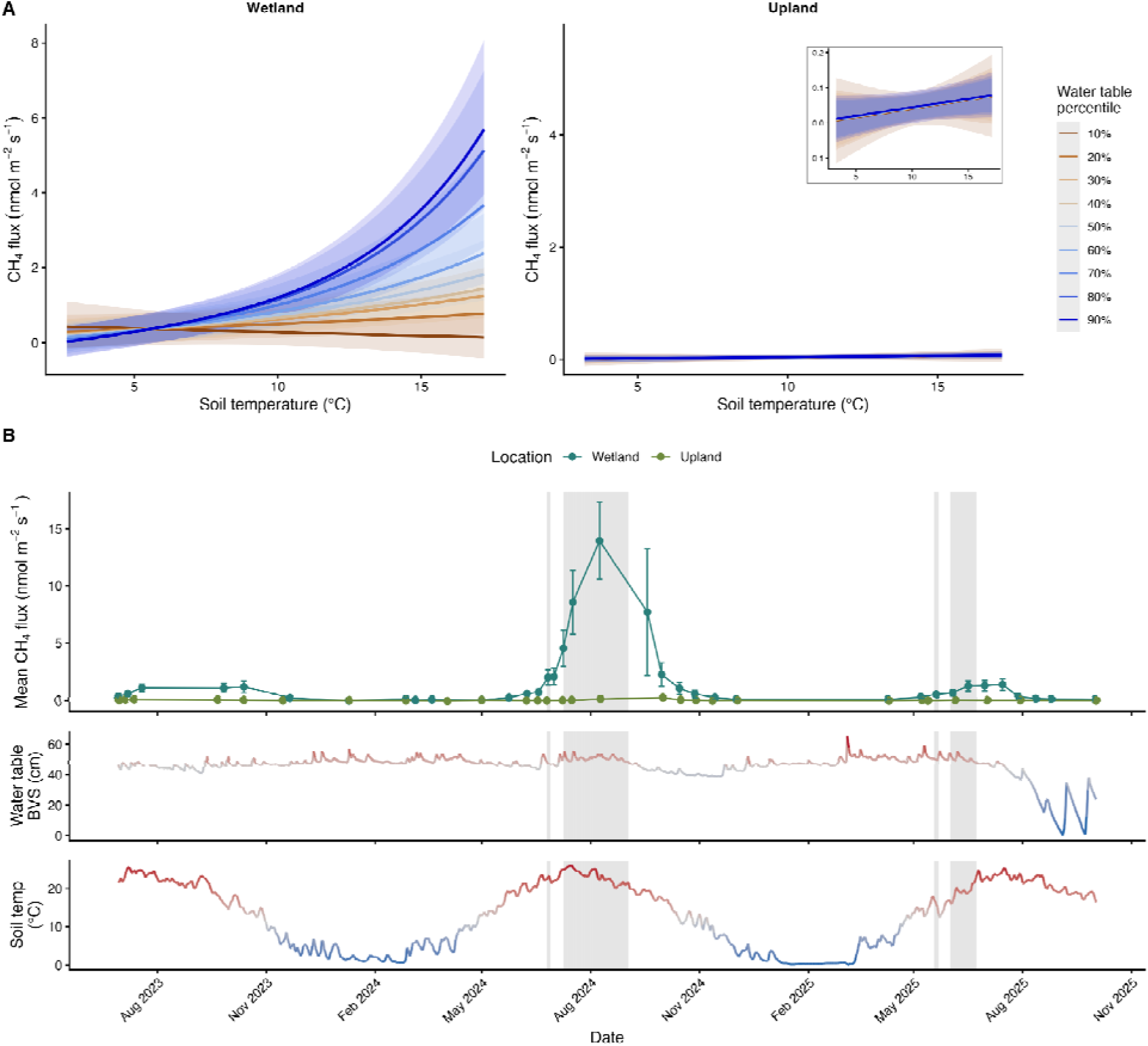
Temperature × water table interactions and temporal correspondence with observed flux. (A) Model predictions (core model: temperature × water table × species) showing CH₄ flux across a temperature gradient at different water table percentiles (color gradient from brown = 10th percentile to dark blue = 90th percentile). Species-averaged predictions; shaded ribbons show 95% confidence intervals. Upland predictions shown on the same y-axis scale as wetland (left) and at expanded scale (inset) to visualize the minimal response. Maximum predicted wetland flux is 72× higher than upland maximum. (B) Temporal pattern of stem CH₄ flux with environmental drivers. Top panel: mean stem CH₄ flux by sampling round at wetland (teal) and upland (olive) sites. Middle and bottom panels: daily water table depth (BVS well) and soil temperature, colored by z-score (blue = below average, red = above average). Gray shading indicates periods when 1-week rolling z-scores for both drivers exceeded 0.5, representing sustained warm and wet conditions.

Species identity contributed substantially to wetland variance, driven primarily by Nyssa sylvatica, which emitted an order of magnitude more CH₄ than co-occurring generalist species. The dominance of N. sylvatica in wetland emissions likely reflects its flood tolerance adaptations. Swamp-adapted Nyssa populations can maintain C supply under flooding (Angelov 1996), and develop morphologically distinct root systems with enlarged air spaces and hypertrophied stem lenticels under prolonged flooding (Hook et al. 1971, Keeley and Franz 1979), creating low-resistance pathways for gas exchange between saturated soils and atmosphere. These conduits, evolved for oxygen transport to roots, may inadvertently facilitate CH₄ efflux, as aerated tissue would not discriminate between oxygen and methane. This interpretation aligns with findings that lenticel density and wood-specific density are key predictors of stem CH₄ emissions across wetland-adapted species (Pangala et al. 2014, 2015), and that bark structural characteristics fundamentally determine CH₄ transport pathways (Jeffrey et al. 2024).

### Uplands: weak environmental control

Environmental drivers poorly predicted upland stem CH₄ flux, with most variation occurring within individual trees over time rather than between species or individuals. There is little evidence for soil production or transport of CH₄ at the upland site. Harvard Forest upland soils are documented CH₄ sinks (Jevon et al.), and upland trees do not typically exhibit the vertical flux gradient characteristic of soil-derived CH₄ transport observed in wetland systems (Barba et al.; Gewirtzman et al.), though we did not measure vertical profiles in this study.

Most upland variability likely reflects competing production and consumption processes (Gewirtzman et al.; Jeffrey et al.; Gauci et al., Leung et al.). When methanogenesis rates approach methanotrophy rates, the balance becomes critical and the net flux more variable and difficult to predict. Tree-specific differences in internal wood moisture, substrate availability, oxygen diffusion rates, and microbial community composition likely create heterogeneity in this production-consumption balance that our environmental measurements could not capture.

Approximately 26.5% of upland observations were negative, indicating net CH₄ uptake, though fluxes were low and sporadic. We were unable to identify environmental controls or tree characteristics explaining when uptake occurred, and negative fluxes were observed across all three upland species. This pattern is consistent with the competing production-consumption framework: when conditions temporarily favor methanotrophic over methanogenic activity, the stem transitions from source to sink. However, further work is needed to confirm that biological methanotrophy is responsible for these observed fluxes.

### Internal wood condition: opposite mechanisms across sites

The relationship between internal wood condition and CH₄ flux was opposite in direction across sites, reflecting fundamentally different CH₄ sources and transport pathways (Figure 5). Specifically, at the upland site, ERT CV was positively correlated with flux, driven by Q. rubra and A. rubrum. Previous work has identified wetwood and decay as potential origins of tree-stem CH₄ production (Zeikus and Ward 1974; Covey et al. 2012; Gewirtzman et al. 2025), and these results support that hypothesis. Trees with moderate decay (Categories II–III) appear most efficient at producing CH₄. Presumably, in these trees, moisture accumulates within partially decayed wood, carbon substrates remain available, and the still-intact wood structure prevents oxygen ingress. Anaerobic microsites then develop where microbes can both draw down oxygen and produce substrates for methanogenesis, particularly hydrogen for hydrogenotrophic methanogenesis, supporting previous observations of wood damage and decay enhancing methane egress (Gorgolewski et al. 2022; Hietala et al. 2015; Marra et al. 2018). We note it is also possible that there is some trace contribution of methane produced by decay fungi themselves (Huang et al. 2022; Lenhart et al. 2012), or of transported methane evading through preferential flowpaths opened by structural decay that our measurements would have missed.

At the wetland site, by contrast, N. sylvatica showed a significant negative correlation between ERT CV and flux. Trees classified as Category I (intact) were the most efficient at transporting soil-derived CH₄. While decay may support in situ methanogenenic conditions as moisture accumulates (Skaar 1988; Brischke and Alfredsen 2020) and microbes draw down oxygen within decaying wood (Röllig et al. 2025), this production is likely trivial compared to the abundant soil-derived CH₄ source. Meanwhile, the increased pore space and moisture content associated with decay are expected to reduce diffusivity, and thus gas transport, through wood (Sorz and Hietz 2006; Thybring 2017). Thus, our data are consistent with the idea that the net effect of decay in wetland trees is reduced transport efficiency, overwhelming any additional decay-associated gross production.

Trees with extensive structural decay (Category IV) may represent a transition state at both sites. When decay progresses to cavity formation and structural loss, oxygen can more readily enter the stem, creating conditions that inhibit strict anaerobes and favor aerobic methanotrophs. If positioned above the water table where wood is less saturated, such trees may begin to function more like upland soil, potentially serving as net CH₄ sinks (Covey et al. 2016).

Figure 7 synthesizes these contrasting mechanisms. In wetland systems with saturated, anoxic soils producing abundant CH₄, healthy trees with intact vascular systems transport soil-derived gas efficiently, with maximum emissions at the stem-atmosphere interface near the water table where concentration gradients are steepest. Internal decay reduces transport efficiency despite supporting minor additional decay-associated production, resulting in lower net emissions from decayed wetland trees. In upland systems with aerobic soils producing minimal CH₄, healthy trees emit only trace amounts. Decay creates anaerobic microsites where in situ methanogenesis can occur, modestly increasing emissions from a near-zero baseline. The opposite relationships between wood decay and flux across sites reflect these fundamentally different sources and transport pathways.

**Figure 7.**
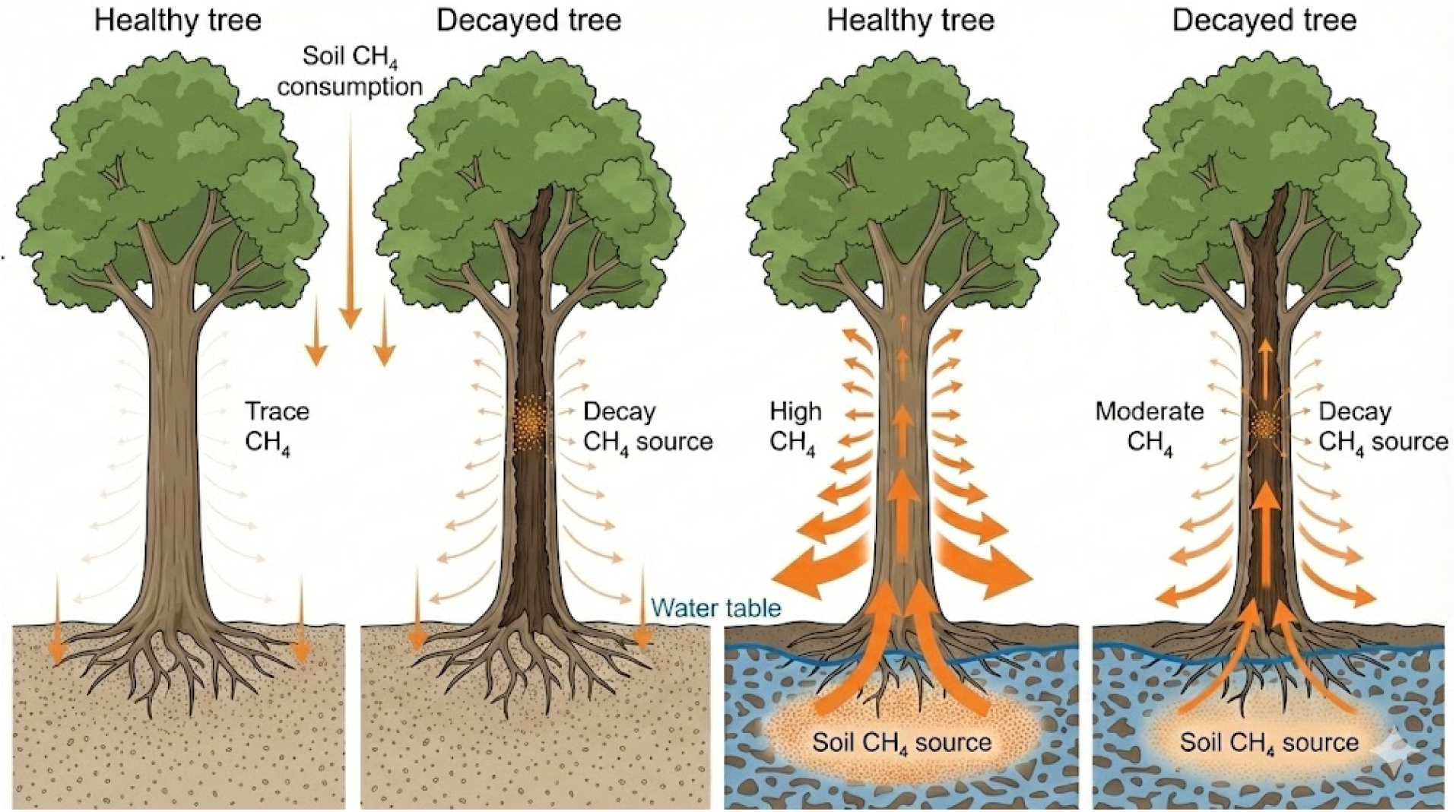
Conceptual model of stem CH₄ flux mechanisms across hydrologic gradients and tree health conditions. Arrows represent CH₄ flux magnitude and direction (arrow thickness proportional to flux). Upland sites (left) have aerobic CH₄-consuming soils with trace CH₄ production in tree stems; decay increases emissions through enhanced in-situ methanogenesis. Wetland sites (right) have saturated soils producing abundant CH₄; healthy trees transport soil-derived CH₄ with maximum emissions at the stem-atmosphere interface near the water table, while decay reduces transport efficiency and net emission despite additional minor decay-associated gross production.

### Implications and future directions

The contrasting variance structures have important implications for scaling and prediction. Wetland flux estimates require accounting for species composition and the presence of high-emitting individuals, particularly N. sylvatica, as a few trees drive stand-level totals, consistent with known hot spot and hot moment control of soil methane emissions (Matthes et al. 2014; Rey-Sanchez et al. 2022). Upland flux estimates are inherently more uncertain due to the large unexplained temporal component; explaining this variation may require measurements at finer temporal resolution or additional mechanistic understanding of methanogen-methanotroph dynamics.

For global CH₄ budgets, these results reinforce that wetland trees are hot spots for forest CH₄ emissions while upland contributions are much smaller per unit area. However, the massive within-wetland heterogeneity means that simple upscaling approaches, multiplying mean flux by wetland forest area, will produce highly uncertain estimates. Accounting for spatial heterogeneity in species composition, water table dynamics, and tree health is essential for accurate landscape-level projections. Meanwhile, upland tree fluxes are at least an order of magnitude lower but nonzero, which can still sum to substantial global fluxes when scaled to the world’s upland forest area. Determining the balance between tree uptake and emission, and the factors controlling that balance (Gauci et al. 2024; Mochidome et al. 2025; Gewirtzman et al. 2025), remains an urgent priority.

Several knowledge gaps warrant further investigation. Our measurements captured net fluxes but could not partition gross production and consumption; isotopic approaches or inhibitor experiments could resolve whether upland stem emissions represent in situ methanogenesis or transport of trace soil-derived CH₄ (von Fischer and Hedin 2007). High-resolution temporal monitoring at wetland sites would clarify the role of short-acting drivers we could not fully assess, such as transpiration effects that covary with energy and water balance variables (Barba et al. 2019). Characterizing the microbial communities within and on tree stems, particularly the relative abundance and activity of methanogens versus methanotrophs, would provide mechanistic insight into the production-consumption balance and the potential for bark-associated CH₄ oxidation (Yang and Silver 2016; Bechtold et al. 2025; Krumholz et al. 1995; Arnold et al. 2023). Understanding variation among wetland tree species in transport capacity, including which species develop aerenchyma and what anatomical or physiological traits distinguish high transporters, would improve predictions of how vegetation composition influences landscape-scale emissions (Bhullar et al. 2013; Anttila et al. 2024), particularly given emerging evidence that vegetation may alter the timing rather than magnitude of wetland CH₄ release (Bansal et al. 2020). The role of plant-soil interactions also deserves attention: does rhizospheric carbon supply from certain species enhance soil methanogenesis (Whiting and Chanton 1993; Määttä and Malhotra 2024), or do species differences primarily reflect morphological transport capacity and microtopographical position? Finally, as extreme precipitation events become more frequent in the northeastern United States and alter wetland hydrology, seasonal water balance, and species composition (Shoemaker et al. 2014; J. H. Matthes et al. 2025; Desprez et al. 2014), understanding how transitional ecosystems shift between source and sink behavior will be essential for predicting landscape-level CH₄ dynamics.

## Conclusions

This study reveals that the controls on tree-mediated methane fluxes differ fundamentally between wetland and upland forests. Wetland trees emitted approximately 40-fold more CH₄ than upland trees, with fluxes strongly regulated by temperature and water table interactions that explained nearly two-thirds of variance, consistent with soil-derived CH₄ transport through tree stems. Species identity was equally important, as the wetland specialist Nyssa sylvatica emitted an order of magnitude more CH₄ than co-occurring generalist species, likely due to flood-tolerance adaptations that create efficient gas transport pathways. In contrast, upland fluxes showed minimal environmental control, with most variance occurring as unexplained temporal variation within individual trees, suggesting competing methanogenic and methanotrophic processes operating near equilibrium. Internal wood condition had opposite effects across sites: decay increased emissions in upland trees by creating anaerobic microsites for in situ methanogenesis, while decay decreased emissions in wetland trees by reducing transport efficiency of the dominant soil-derived CH₄ source. These findings demonstrate that accurate landscape-scale CH₄ budgets require accounting for species composition, hydrologic setting, and tree health, and highlight the need for further research into the microbial and physiological mechanisms underlying the production-consumption balance in upland forest trees.

## Acknowledgements

We thank Audrey Barker-Plotkin, Annabelle Rayson, Ashley Dawn, Sonia McCollum, Charles Harvey, Missy Holbrook, the Harvard Forest Woods Crew, and the Harvard Forest Summer Research Program in Ecology staff for field and logistical support. Mark Bradford and Peter Raymond provided guidance throughout this project.

J.G. was supported by a National Science Foundation Graduate Research Fellowship (DGE-2139841), the Yale Institute for Biospheric Studies, the Kohlberg-Donohoe Research Fellowship, and a Harvard Forest LTER Graduate Student Research Award. This research was supported by National Science Foundation grant DEB-194592 and Department of Energy Environmental System Science grant DE-SC0024092 to J.H.M..

Environmental data were provided by the Harvard Forest Long-Term Ecological Research program (NSF grants DEB-8811764, DEB-9411975, DEB-0080592, DEB-0620443, DEB-1237491, DEB-1832210). Data used in this research were provided by the PhenoCam Network, which has been supported by the National Science Foundation, the Long-Term Agroecosystem Research (LTAR) network which is supported by the United States Department of Agriculture (USDA), the U.S. Department of Energy, the U.S. Geological Survey, the Northeastern States Research Cooperative, and the USA National Phenology Network. We thank the PhenoCam Network collaborators, including site PIs and technicians, for publicly sharing the data that were used in this paper. Soil water content data were provided by the National Ecological Observatory Network, a program sponsored by the National Science Foundation and operated under cooperative agreement by Battelle.

## Supporting Information

**SI Table 1.** Wetland model comparison showing predictor selection and model fit.

| Model | Predictors | R<sup>2</sup> (%) | AIC | BIC | Nparam |
| --- | --- | --- | --- | --- | --- |
| Core | Temp × WTD × Species | 65.4 | 831.0 | 889.9 | 12 |
| Full | Core + LE × Species + SWC × Species | 67.6 | 784.8 | 868.9 | 18 |
| Alt: SWC core | Temp × SWC × Species | 61.7 | 914.9 | 973.7 | 12 |
| Alt: LE core | LE × WTD × Species | 56.5 | 1016.0 | 1074.9 | 12 |

*Note: Temp = soil temperature at Ha2 (282h window); WTD = water table depth at BVS well (132h window); LE = latent heat flux at Ha1 (123h window); SWC = soil water content from NEON (129h window). All predictors standardized. N = 495 observations, 30 trees.*

**SI Table 2.** Species-specific environmental effects (standardized coefficients) for wetland models.

| **Species** | **Model** | **Temperature** | **Water table** | **Temp × WTD** |
| --- | --- | --- | --- | --- |
| *N. sylvatica* | Core | 0.552*** | 0.534*** | 0.415*** |
|  | Full | 0.779*** | 0.927*** | 0.422*** |
| *T. canadensis* | Core | 0.078*** | 0.074*** | 0.038*** |
|  | Full | 0.025*** | 0.154*** | 0.084*** |
| *A. rubrum* | Core | 0.111*** | 0.190*** | 0.165** |
|  | Full | 0.152*** | 0.254*** | 0.163** |

\*\*\* $p < 0.001$ ; \*\* $p < 0.01$ ; \* $p < 0.05$ . Core model: temperature (282h) × water table (132h) × species. Full model adds latent heat (123h) × species and soil water content (129h) × species.

**SI Table 3.** Upland model comparison.

| **Model** | **Predictors** | **Window** | **R<sup>2</sup> (%)** | **AIC** | **BIC** |
| --- | --- | --- | --- | --- | --- |
| Model A | TS_Ha1 × SWC × Species | 1h | 8.8 | −398.1 | −337.5 |
| Model B | TS_Ha2 × WTD × Species | 282h/132h | 8.2 | −455.0 | −394.1 |

*Note: TS = soil temperature; SWC = soil water content (NEON); WTD = water table depth (BVS well). Model B uses identical predictors and windows as wetland model for direct comparison. N = 570 observations, 30 trees.*

**SI Table 4.** Cross-site comparison of species-specific environmental responses (standardized coefficients).

| Site | Species | Temperature | Moisture | Temp × Moisture |
| --- | --- | --- | --- | --- |
| **Wetland** | *N. sylvatica* | 0.552*** | 0.534*** | 0.415*** |
|  | *T. canadensis* | 0.078*** | 0.074*** | 0.038*** |
|  | *A. rubrum* | 0.111*** | 0.190*** | 0.165** |
| **Upland** | *Q. rubra* | 0.015 | −0.005 | −0.014 |
|  | *T. canadensis* | −0.003 | 0.019 | 0.023 |
|  | *A. rubrum* | 0.056** | −0.010 | −0.009 |

\*\*\* $p < 0.001$ ; \*\* $p < 0.01$ ; \* $p < 0.05$ . Wetland moisture = water table depth (132h window); Upland moisture = water table depth (132h window) from Model B for direct comparison. Both models use soil temperature with matched windows (282h). Wetland: Core model, $N = 495$ ; Upland: Model B, $N = 570$ .

